# Transfer learning from simulations improves the classification of OCT images of glandular epithelia

**DOI:** 10.1101/2020.10.26.355180

**Authors:** Sassan Ostvar, Han Truong, Elisabeth R. Silver, Charles J. Lightdale, Chin Hur, Nicholas P. Tatonetti

## Abstract

Esophageal adenocarcinoma (EAC) is a rare but lethal cancer with rising incidence in several global hotspots including the United States. The five-year survival rate for patients diagnosed with advanced disease can be as low as 5% in EAC, making early detection and preventive intervention crucial. The current standard of care for EAC targets patients with Barrett’s esophagus (BE), the main precursor to EAC and a relatively common condition in adults with chronic acid reflux disease. Preventive care for EAC requires repeated surveillance endoscopies of BE patients with biopsy sampling, and can be intrusive, error-prone, and costly. The integration of minimally-invasive subsurface tissue imaging in the current standard of care can reduce the need for exhaustive tissue sampling and improve the quality of life in BE patients. Effective adoption of subsurface imaging in EAC care can be facilitated by computer-aided detection (CAD) systems based on deep learning. Despite their recent successes in lung and breast cancer imaging, the development of deep neural networks for rare conditions like EAC remains challenging due to data scarcity, heavy bias in existing datasets toward non-cases, and uncertainty in image labels. Here we explore the use of synthetic datasets–specifically data derived from simulations of optical back-scattering during imaging– in the development of CAD systems based on deep learning. As a proof of concept, we studied the binary classification of esophageal OCT into normal squamous and glandular mucosae, typical of BE. We found that deep convolutional networks trained on synthetic data had improved performance over models trained on clinical datasets with uncertain labels. Model performance also improved with dataset size during training on synthetic data. Our findings demonstrate the utility of transfer from simulations to real data in the context of medical imaging, especially in the severely data-poor regime and when significant uncertainty in labels are present, and motivate further development of transfer learning from simulations to aid the development of CAD for rare malignancies.

## I. Introduction

Esophageal adenocarcinoma (EAC), a cancer of the distal esophagus, is a public health concern in several countries including the United States due to its quickly rising incidence and poor prognosis. The current 5-year survival rate for EAC is ~20% in the US and drops to <5% for patients diagnosed with late-stage disease, calling to attention the need for improved preventive screening of at-risk patients [1]. Population surveillance for EAC targets Barrett’s esophagus (BE), or pre-malignant intestinal metaplasia of the distal esophageal mucosa. BE affects an estimated 2-5% of the US adult population [2], a small fraction of whom develop cancer. Preventive screening for EAC is achieved by repeated surveillance endoscopies that rely on a combination of visual examination, mucosal biopsies, and endomicroscopy [1], [3]. Despite efforts to optimize EAC surveillance for early detection, malignant progressions in BE (i.e. dysplasia) remain difficult to detect with existing practices, and a balance between diagnostic yield, cost-effectiveness, and intrusiveness of screening is yet to be reached. Advanced imaging modalities like optical coherence tomography (OCT) are emerging technologies that may improve the accuracy and reduce the intrusiveness of the standard of care in endoscopic EAC surveillance [4], [5]. Subsurface tissue imaging with OCT provides rich depth-resolved structural information on the entire distal esophagus. However, the difficulty associated with interpreting this data under time constraints is a barrier to its effective integration with existing procedures, especially by non-expert endoscopists outside specialized care centers.

Deep learning (DL) for computer-aided detection (CAD) has recently led to breakthroughs for similar surveillance targets in screening mammography for breast cancer [6], low-dose chest CT for lung cancer [7], and OCT-based diagnostics in ophthalmology [8], [9], often achieving similar performance to human raters [10]. Artificial intelligence (AI) systems for CAD can improve cancer diagnostics by increasing the accuracy of image-based early detection, reducing the required human workload in surveillance programs, and minimizing the morbidity of preventive interventions [6]. Advances in early detection can directly impact patient outcomes and improve the effectiveness and cost-effectiveness of population surveillance. The potential benefits are similarly multi-fold to the standard of care for BE, where the likelihood of missed malignancies, over-screening, and propensity for risk-averse but aggressive interventions like complete eradication of the affected esophageal mucosa are currently problematic. Similar to many other examples in medical imaging, the application of DL to OCT imaging of the esophagus for cancer surveillance is limited by data scarcity–that is, the difficulty of curating sizable datasets with reliable and balanced labels [11].

Transfer learning (i.e. the use of pre-trained model architectures to develop image classifiers [12]), has recently been adopted to overcome data scarcity limitations in medical imaging for applications in radiology [13], ophthalmology [14], and brain imaging [15], [16]. The accessibility of computer vision benchmarking datasets such as ImageNet have made them a popular choice for pre-training, but it is not clear if transfer from natural to medical datasets is optimal in deep convolutional architectures [17]. Medical imaging datasets are generated by measurements for specific materials over precisely-selected bands of the electromagnetic spectrum and under controlled experimental conditions. Benchmarking datasets instead tend to span many scales, materials, light sources, and devices, and cluster around the visible spectrum. Another point of divergence is the existence of thousands versus a handful of labels in the classification problems defined for the two types of data.

Here, we make an argument in favor of fine-tuning DL models on synthetic datasets derived from simulations of the imaging process. Such a dataset can be constructed based on knowledge of tissue composition and structure in cases and controls, subsurface light scattering, and the process of signal construction in an imaging method of interest. We explore this idea using an esophagus OCT dataset collected at the Columbia University Irving Medical Center between 2014 to 2018. We focus on BE as the first structural transition along the EAC pathway, where the stratified epithelium of the healthy esophagus is replaced with a glandular epithelium that mimics the gastric and intestinal morphologies. Microscopic examination of this lesion reveals a complete restructuring of the affected tissue into a glandular mucosa, resulting in a loss of clear lamination between the epithelium and the stroma, which is reflected in the OCT signal [5].

We frame our analysis as binary classification of OCT images into metaplastic and normal segments using instances of the ResNet-18 architecture starting with ImageNet weights. Model performance was evaluated after fine-tuning on (i) synthetic data and (ii) clinical data with noisy annotations inferred from electronic health records (no retrospective expert annotations). Both models were evaluated using an external validation set with retrospective expert annotations. Fine-tuning on the synthetic dataset led to significant improvement in performance above chance. In comparison, fine-tuning on the clinical dataset with noisy annotations led to marginal improvement over chance. We discuss these results in the context of automatic segmentation of esophagus OCT into normal and metaplastic regions. Finally, the prospects for improving the proposed pipeline by increasing the fidelity of physics-based data synthesis are briefly discussed as a template for future work. We argue that transfer learning from simulations enables the integration of knowledge of disease from disparate sources, modalities, and scales, and improve model development for CAD in data-poor settings.

## II. Methods

### A. Clinical dataset

#### 1) Patients

We identified a retrospective cross-sectional cohort of patients with both (i) confirmed diagnosis of BE, and (ii) at least one upper endoscopy encounter with OCT imaging at the Columbia University Irving Medical Center (CUIMC). For each patient we retrieved volumetric OCT scans of the distal esophagus and the associated hospital electronic health records (EHR), including diagnoses and treatment histories, gastrointestinal endoscopy reports, and the pathology reports describing biospecimens that were collected and analyzed during the course of surveillance and treatment. We carried out all data curation and management according to the rules set by a CUIMC Institutional Review Board to ensure the privacy of human subjects and fair use of data.

#### 2) OCT scans

We obtained 3D OCT scans and associated DICOM metadata for each encounter directly from a swept-source instrument (NvisionVLE^®^ Imaging System, NinePoint Medical, Bedford, MA) in collaboration with the Division of Digestive and Liver Diseases at CUIMC. We identified three types of scans: *full scans*, which covered the entire tissue segment without manual guidance, and *manual scans*, which covered areas of interest to the endoscopist, and preparatory ‘scout’ scans. Each volumetric scan typically resolved part of the stomach or a hiatial hernia below the GE junction, to an extent that varied case by case. Full scans contained 1200 cross-sectional B-scans (hereafter ‘frames’), and the stack height varied for manual scans. Each frame was a 2048 × 4096 image recorded in 8-bit grayscale. We analyzed the resulting dataset on the basis of subdivisions of frames as described below. The instrument employed balloon catheters to dilate and immobilize the esophageal wall during image acquisition. The balloon diameters and operative pressures showed variation over the period of data acquisition (Fig. 3, Supplementary Information). In each frame, the balloon cross-section was discernible as a line of bright pixel intensity marking the boundary between the tissue surface and the esophageal lumen. The balloon catheter also provided a (longitudinal) registration watermark that served as the reference for the measurement of circumferential positions. We normalized all 2048 × 4096 frames prior to the analysis to flatten the epithelial surface and remove the (dark) lumen using the balloon cross-section pixel intensity as the threshold.

#### 3) Annotations derived from EHR

We thoroughly examined the pathology reports with readings describing biospecimens to deduce a set of corresponding labels and sampling locations for each tissue sample. To generate the final labels, we successively reduced an exhaustive dictionary of terms that had been used in the reports to describe the biospecimens (Table II, Supplementary Information). This process resulted in four primary clinical phenotypes of interest: `NORMAL` indicated an absence of evidence supporting a diagnosis of intestinal metaplasia, dysplasia, or cancer (i.e. the stratified epithelium was preserved). `METAPLASIA` indicated endoscopic and pathology findings supporting a diagnosis of Barrett’s metaplasia. `DYSPLASIA` indicated pathology findings indicating disease progression in the form of glandular dysplasia. `CANCER` indicated observations of malignant neoplasia of any clinical stage, including intramucosal carcinomas, cancers with submucosal invasion, etc. Additionally, we marked stomach tissue samples under `STOMACH` and all other under `OTHER`. Tissue sampling locations had been reported in GI endoscopy reports as pairs of longitudinal (‘distance to incisors’) and angular (‘clock’) positions. We asked two raters to independently match the pathology readings with ROIs in the scans by first matching the entries in pathology reports with the pair of longitudinal and angular values provided in the GI endoscopy reports, and then converting the recordings to approximate regions of interest (ROIs) in the scan’s coordinate system. For a subset of patients, a laser marking device had been used to mark the precise location of the tissue sample, enabling improved matching of records with ROIs.

#### 4) External validation set

We evaluated the performance of image classifiers using an independently annotated set of frames that was prepared retrospectively in collaboration with an expert gastroenterologist and frequent user of the OCT instrument (CJL). To generate this set, we asked the rater to assign annotations and a confidence score between 50-100% to the regions within a set of pre-selected frames that were suspected for `METAPLASIA` (cases) or `NORMAL` (controls). We then subdivided this set of annotated frames into patches of 128×256 pixels and used them as the primary benchmark for model performance evaluation.

### B. Image classification with deep convolutional networks

We performed image classification experiments on **128** × **256** 8-bit grayscale patches using instances of the ResNet-18 architecture starting with ImageNet weights [18]. We evaluated all model instances after 40 epochs of training with a learning rate of 3 ×**10**^-6^, learning rate decay over 7 epochs of 0.5, and batch size of 600, and repeated each run with 100 model instances that were identical except in the last fully connected layer, which we initiazed randomly for each run. In our main analysis, we compared model performance after training on two independent datasets: (i) subsets of the clinical dataset with annotations deduced from pathology reports, and (ii) a synthetic dataset derived from simulations, both prepared using a 70/30 training/testing split. We measured model performance using the area under the curve of the receiver operating characteristic curve (ROC AUC). In this study, we report the results of binary supervised classification of patches into `METAPLASIA` (cases) or `NORMAL` (controls).

### C. Synthetic data

**Fig. 1.**
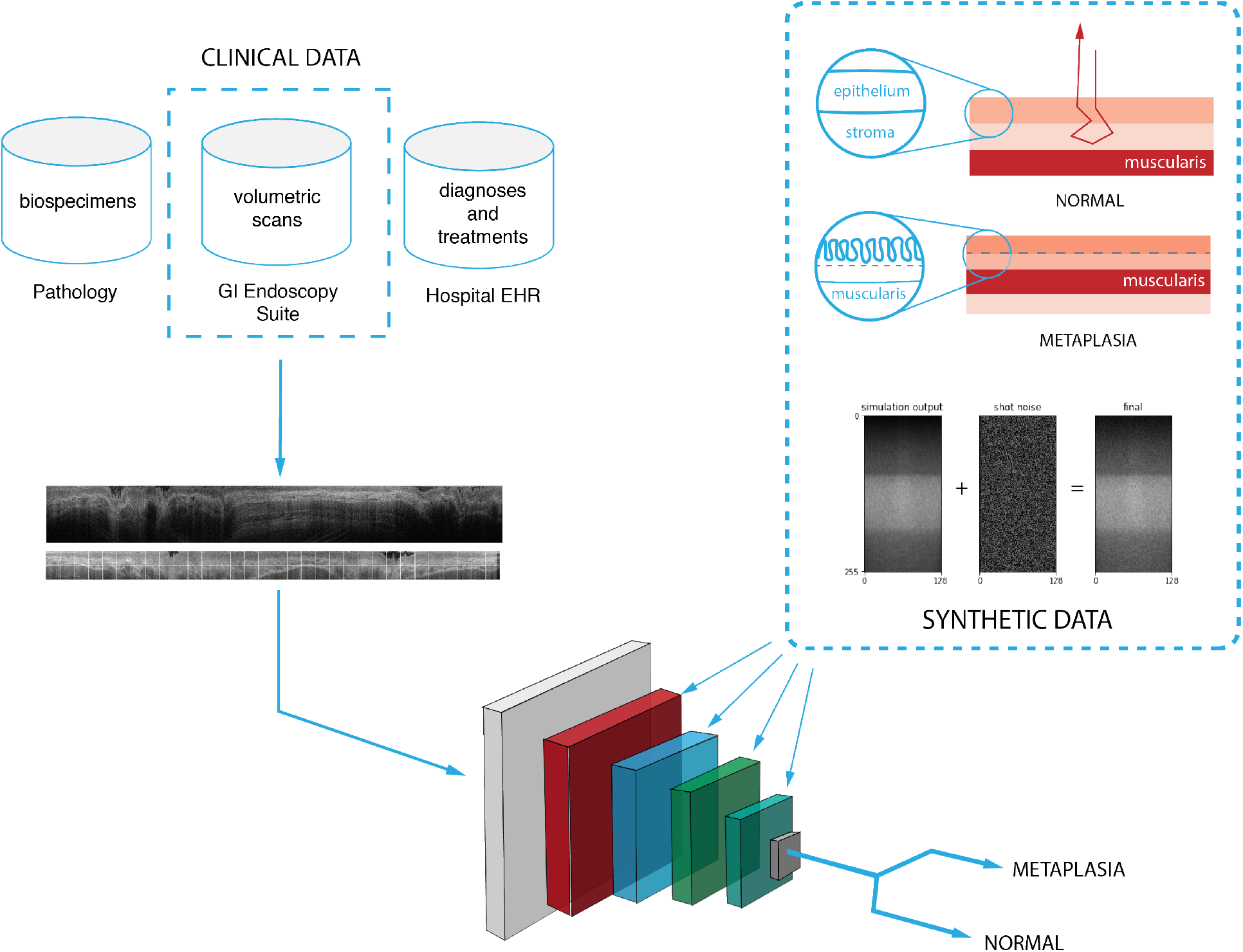
Overview of the model development pipeline: Instances of ResNet-18 with ImageNet weights are trained on synthetic data derived from simulations of light scattering in model tissue geometries. The resulting model is used to classify clinical OCT data into patches indicating either a normal squamous epithelium (`NORMAL`) or Barrett’s esophagus (`METAPLASIA`).

Subsurface imaging for EAC surveillance targets the esophageal mucosa (EM). The EM is a multilayered structure consisting of a stratified epithelium, stroma, and a muscularis layer. A clear stratification between the epithelium and stroma is present in `NORMAL` and partially lost in `METAPLASIA`. In the following, we outline a set of simulation experiments conducted to approximate patches of `NORMAL` and `METAPLASIA` in our dataset on basis of this structural difference using simple multilayered representations of the EM. Clinical OCT frames are constructed out of groups of adjacent axial optical reflectivity profiles (A-lines). Each OCT frame encodes structural information as variations in optical reflectivity in the axial and transverse directions. Simulation of OCT data can therefore be broadly considered as the problem of computing a series of A-lines for given spatial distributions of tissue constituents. Simulation of OCT A-lines is the focus of computational optical imaging (COI), a research program dedicated to the simulation of light scattering in biological tissue and signal localization in imaging instruments based on first principles or approximate sampling methods as described in the following (e.g. [19], [20]).

#### 1) OCT data structure and artifacts

Each OCT scan of the lower esophagus is an *n_r_* × *n_θ_* × *n_z_* matrix ***R*** = *R_ijk_* of voxel intensities that discretizes the annulus bound by *r*_0_ ≤ *r* ≤ *r*_0_ + *n_r_***Δ***ℓ_r_* and 0 ≤ *z* ≤ *n_z_* **Δ***ℓ_z_* with voxel resolutions **Δ***ℓ_voxel_* = (**Δ***ℓ_r_*, **Δ***ℓ_θ_*, **Δ***ℓ_z_*), where *r*_0_ is the radius of the inflated balloon catheter. A volumetric scan is interpreted as a stack of *n_z_* frames (or B-scans) ***R**_k_* = *R_ij,k_*, which are in turn composites of *n_θ_* axial reflectivity profiles (A-lines) ***R**_kj_* = *R_kj,i_*. The instrument records ***R*** one A-line at a time during a helical pull-back of the endoscopic probe with two degrees of freedom that control the longitudinal (**Δ***ℓ_z_*) and transverse (**Δ***ℓ_θ_*) voxel resolutions, and the coherent length of the light source sets the axial resolution (**Δ***ℓ_r_*). We assume that *n_r_, n_θ_*, and *n_z_* are constant across experiments. Ideally, the instrument’s light source and fiber optic probe coincide with the centroid of the catheter’s cross-section during image acquisition, but a persistent offset is often present in practice, which may affect the total imaging depth. Each frame is susceptible to motion artifacts due to in-plane displacements of the endoscopic probe at fixed z. The final image also carries shot noise that is introduced as the signal is transmitted through the fiber optic probe and the optical detector’s circuitry.

#### 2) Simulated optical back-scattering

To estimate the A-lines, we adopted a mesh-based Monte Carlo (MC) algorithm to simulate subsurface scattering in model tissue geometries with predefined optical properties [21]–[23]. The MC method provides an estimate of the spatial distribution of the energy of back-scattered radiation via sampling a set of possible trajectories of individual ‘photon packets’ as they interact with mesh elements. Photons are launched from a source and collected by a probe that employs a threshold on the incidence angle of back-scattered packets. Each recorded packet is specified by an optical depth *ℓ_n_* (eqv. to one half the optical path length) and a dimensionless measure of energy, *w_n_* (weight). Packet trajectories are determined by three types of mesh-photon interactions: (i) specular (Fresnel) reflection at the mesh surface, (ii) partial loss of energy proportional to a local adsorption coefficient, *μ_a_*, and (iii) scattering. The former is calculated from the differences in the real part of the refractive index n [20]. The latter is specified by the scattering coefficient, *μ_s_*, and a scattering phase function, *p*(*s′* → *s*) = *p*(*θ, φ*), i.e. the probability density of scattering into a direction *s* given current direction *s′*, parametrized over the polar and azimuthal scattering angles *θ* and *φ* at the site of scatter. In biomedical optics, the dependence of *p*(*θ, φ*) on *φ* and *θ* are typically approximated by the continuous uniform and the Henyey-Greenstein (HG) probability density functions, respectively [24]. The HG function is adjusted by a single variable, −1 ≤ *g* ≤ 1, i.e. the local anisotropy, where −1 and +1 specify dominant backscattering and forward scattering, respectively.

#### 3) Signal localization

Part of the back-scattered radiation that is incident on the instrument’s fiber optic probe is collected and converted into an electric current by an optical detector, from which A-lines are derived. Considering transport in a coordinate system where tissue depth is parameterized by r, this process can be simulated using an indicator function *I*(*r,n*) that enforces the probe’s radial and angular thresholds on the n-th back-reflected packet, and discretizes the axial span with a resolution set by the coherence length *ℓ_c_* of the light source [22]:

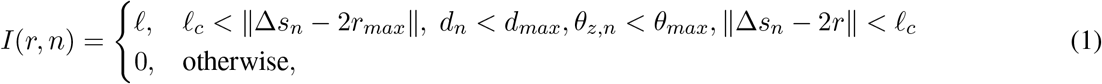

where *d_max_* and *θ_max_* are the positional and angular thresholds of the probe. In Eq-1, *d_n_* is measures the distance between the probe and a reflected packet, **Δ***s_n_* = 2*ℓ_n_* measures the optical path of the n-th back-reflected packet, *θ_z,n_* is the angle of the packet’s trajectory with respect to the r-axis of the lab reference frame, and *r_max_* is the maximum depth reached by the photon packet. The depth-resolved reflectance is then calculated as

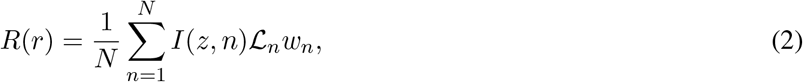

The correction factor 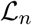 is a likelihood ratio that compensates for biased scattering in the calculation of *ℓ_n_* and *w_n_*. We introduced biased scattering artificially as discussed in [22], [24], [25] to speed up the calculation of R, since most tissue materials have anisotropies close to unity (i.e. dominantly forward scattering), requiring a prohibitively large number of packets to be simulated in order to generate a reliable signal. The signal associated with Eqs 1 and 2 is an estimate of the back-reflected power distribution along the tissue depth, contributed by multiply scattered packets. A similar procedure gives the contribution of ballistic and semi-ballistic back-reflection events [22].

#### 4) Tissue structure

We treated the esophageal tissue as a composite of epithelial, stromal, and muscularis layers. The optical properties of each layer are specified as four scalar fields, *n* = *n*(*r, θ, z*), *μ_a_ = μ_a_*(*r, θ, z*), *μ_s_ = μ_s_*(*r, θ, z*), and *g* = *g*(*r, θ, z*), which are discretized using a tetrahedral mesh. In the simplest approximation, we can study the esophageal cross-section in the limit of vanishing displacement from a perfectly dilated reference configuration (here idealized as a multilayered annulus). We neglect the curvature of the annulus over segments corresponding to 128×256 patches and assume material homogeneity in each layer. The choice of numeric values for *μ_s_, μ_a_, g*, and *n* is guided by published experimental work on the optical properties of gut mucosa over the spectral window of the OCT instrument (1250–1350 nm) [26]–[29].

### D. Simulations

We implemented all classifiers in PyTorch and trained all models using a local NVIDIA Tesla GPU cluster. We used TetGen [30] to generate 3D tetrahedral meshes and an open-source implementation of the Monte Carlo method in CUDA C [22] to perform the subsurface scattering simulations on an NVIDIA Quadro P6000 card. We implemented all pipelines in Jupyter notebooks.

## III. Results

### A. Patient population

Table I provides a summary of the CUIMC patient population and associated pathology and imaging data. Patients met the inclusions criteria if they had a diagnosis of BE and at least one OCT scan of the lower esophagus. We identified a total of 189 BE patients, among whom 43.9% had progressed to BE dysplasia and 12.7% to cancer during their health history. Hiatial hernias were present in 67.2% of the patients.

**TABLE I.** Summary statistics of the CUIMC patient cohort and associated OCT and pathology datasets.

| Variable | Values |
| --- | --- |
| Data period | 2014-2018 |
| Number of patients | 189 |
| Age | 70.0 ± 9.4 |
| Sex (% male) | 130/188 (69.2) |
| High-grade dysplasia (%) | 83/189 (43.9) |
| Malignant neoplasia (%) | 24/189 (12.7) |
| Family history of GI cancers | 19/189 (10.1) |
| Reflux esophagitis | 49/189 (25.9) |
| Atrophic gastritis | 31/189 (16.4) |
| Hiatial hernia (%) | 127/189 (67.2) |
| Duodenitis (%) | 11/189 (5.8) |
| Number of OCT scans | 508 |
| Full scans (%) | 299/508 (58.9) |
| Manual scans (%) | 147/508 (28.9) |
| Scout (test) scans (%) | 62/508 (12.2) |
| Number of biospecimens | 552 |
| Biopsies (%) | 527/552 (95.5) |
| Endoscopic mucosal resection (%) | 12/552 (2.2) |
| Endoscopic submucosal dissection (%) | 7/552 (1.3) |
| Number of laser markings | 118 |

### B. Imaging dataset and annotations

We obtained a total of 508 scans, of which 299 were full scans, covering an invariant 6 cm of tissue that coincided with the lower esophagus and gastric cardia. We excluded partial scans from the study. We assigned frame-by-frame annotations to each scan based on the indications extracted from pathology reports and the approximate locations of tissue samples in the corresponding scan as reported in the GI endoscopy reports. In deriving the annotations, we extracted and analyzed an exhaustive dictionary of descriptive terms from pathology reports, and reduced them to a small set of labels that indicated the extent of disease progression as described in the Supplementary Information (Table II). Measurement of longitudinal and angular positions of tissue samples were based on readings from the regular endoscopic probe. The large discrepancy in how precisely the regular endoscopic probe and the OCT catheter measured distances limited precise assignment of readings from pathology reports to the corresponding sampling locations in the scans. We therefore restricted the clinical training set to the subset of encounters in which at least one biospecimen had been collected with the aid of laser markings (118 in total). We assigned the corresponding annotation to the frames within a given distance from the laser marking (equal to typical dimensions of biopsies obtained using large-capacity forceps, cold forceps, endoscopic mucosal resection, or endoscopic submucosal dissection as indicated in the EHR). The resulting annotated ranges of frames constituted a training/testing dataset of 128 × 256 patches (Fig. 2-C). Significant variation and bias toward controls was present in the pathology dataset (Fig. 3, Supplementary Information).

**Fig. 2.**
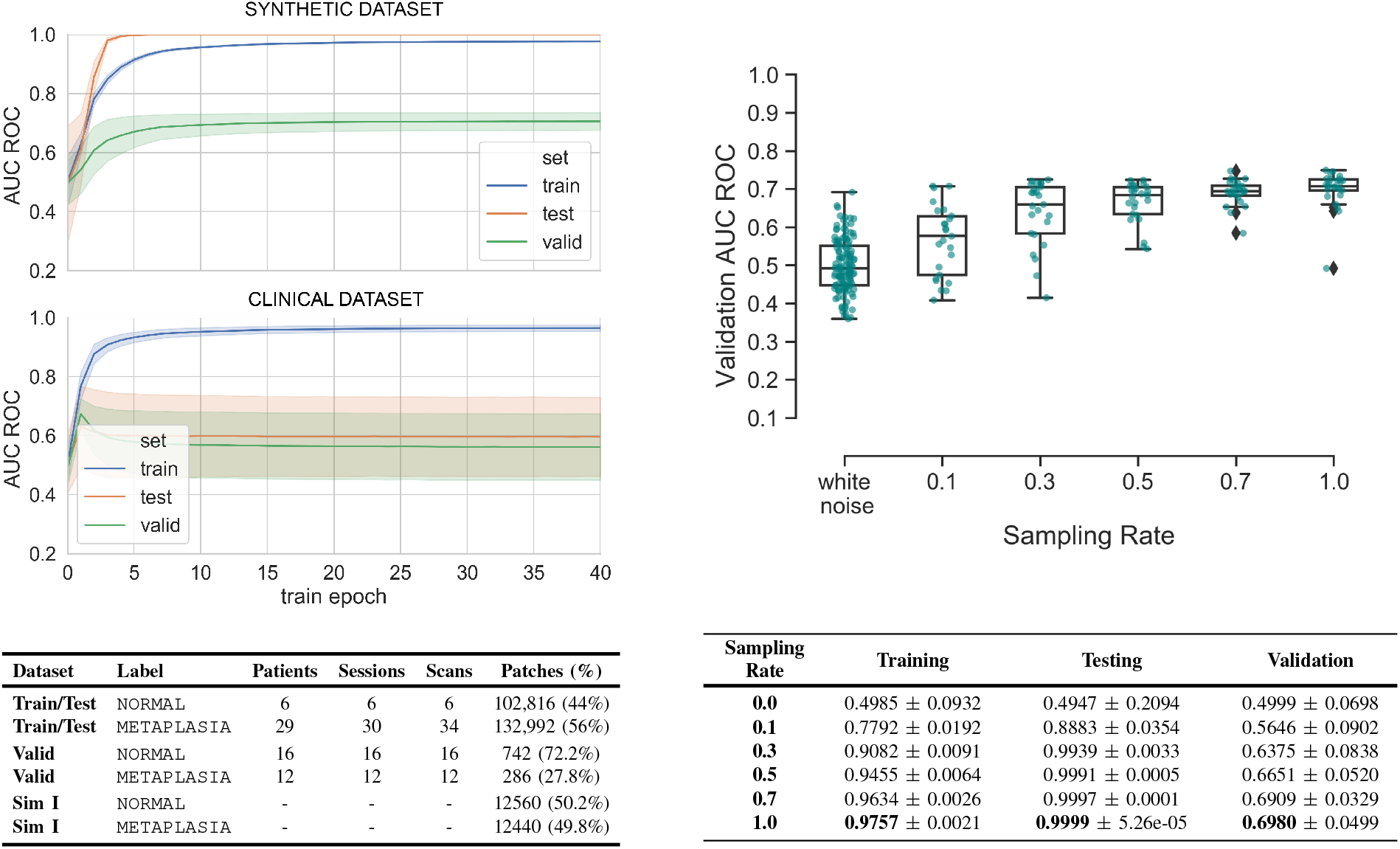
Model performance evaluation after fine-tuning on synthetic and clinical data. (A; top left) Classification performance of ResNet-18 with ImageNet weights over epochs of training on synthetic dataset I, and a clinical dataset with noisy annotations; (B; top right) Classification performance on the validation set as a function of the sampling rate of the synthetic dataset during model training; (C; bottom left) Overview of both training datasets and the validation set; (D; bottom right) Tabulated results corresponding to B.

### C. Simulation of OCT B-scans

We designed the simulated as a set of 128×256 patches equally partitioned between the `METAPLASIA` and `NORMAL` labels. We adopted an idealized model geometry with three distinct layers to simulate each label, and approximated the scattering and adsorption coefficients and anisotropy of each layer based on the literature on optical imaging of gut mucosa in the 1250-1350 nm range of source wavelengths. In setting these parameters, we assumed the muscularis and stromal layers had the same optical properties in both labels, but the epithelial layer in `METAPLASIA` had increased adsorption and scattering coefficients and decreased anisotropy compared with `NORMAL`. We then derived synthetic OCT A-lines from power distributions of back-scattered radiation using a mesh-based Monte Carlo light light scattering algorithm. The algorithm constructed image patches (i.e. subregions of OCT B-scans) one A-line at a time for 128 placements of the light source and probe over the model geometry and clipped to 256 pixels in the axial direction. We augmented the resulting dataset by a factor of 10 by permuting a white noise floor in the range [15, 25] dB and re-scaling the total pixel intensity to [100, 200] dB, assuming a value of 255 corresponded to the reflectance of the balloon catheter material.

### D. Classification accuracy

We first trained a set of ResNet-18 instances with ImageNet weights on a clinical dataset comprised, respectively, of 102,816 and 132,992 patches of `NORMAL` and `METAPLASIA`. We evaluated the classification accuracy of this model on the external validation set and observed generally poor performance over 40 train epochs as illustrated in Fig. 2-A. As a comparator, we trained an independent set of ResNet-18 instances on a synthetic dataset comprised of ~ 12,500 patches per label, also starting with ImageNet weights, and evaluated for classification accuracy using the validation set. Training on the synthetic dataset yielded a peak mean AUC ROC of ≈ 0.7 over 40 train epochs (Fig. 2-A). We then repeated the experiments with synthetic data, this time varying the fraction of data used during model training between 0.0 and 1.0, using a dataset of white noise images as the negative control. Fig. 2-B illustrates the evolution in the mean and variance of the resulting AUC ROC as a function of the sampling rate, where mean performance shows a monotonous increase. Similarly, the variance in performance between different instances shows a decreasing trend.

## IV. Discussion

CAD systems based on models of computer vision employing deep learning rely on sizable and precisely annotated clinical datasets. The curation of such datasets is laborious as it may require extensive retrospective expert evaluation. These datasets may be further limited in size, heavily biased toward cases or controls, and scattered across different institutions for clinical conditions of low population prevalence. In this work, we have reported the use of synthetic imaging data to boost the performance of a deep classifier of epithelial disease in one such data-poor setting. We found that fine-tuning a pre-trained deep convolutional architecture on synthetic data derived from simulations of light scattering provided a performance advantage to fine-tuning on a (larger) clinical dataset with noisy labels. To the best of our knowledge, this work is the first to demonstrate transfer from simulated to clinical data in the context of biomedical imaging.

In this study, the performance obtained from the model trained on clinical data was poorer than that obtained from the model trained on simulated data. We speculate that this discrepancy can be explained by persistent uncertainty in the labels of regions within a scanned volume. We have identified several contributing factors to this uncertainty, including limited spatial coverage of biopsy sampling, variation in biospecimen size, and imprecise measurement of longitudinal and angular positions corresponding to sampled regions within a scan. We expect the issues encountered here to be typical of similar datasets in other rare cancers and diseases. Transfer learning from simulations can be regarded as leveraging computation in a directed manner to address both (i) imprecise annotations, and (ii) imbalanced datasets, operating at a trade-off between fidelity and computational tractability. When scalable computations with properly motivated models are possible, they may reduce the burden of retrospective data surveys and enable the development of otherwise unreliable classifiers.

We can expand the simulations performed in this study in a number of ways. Accounting for residual stress and thickness inhomogeneities [31] and deformations induced by the balloon catheter [32], [33] can improve the modeling of tissue configuration during imaging. Simulations of wave scattering and signal localization in OCT can be based on first-principles calculations using Maxwell’s equations, although this method is currently computationally prohibitive [19], [34], [35]. Explicitly accounting for light source geometry and signal localization in frequency-domain OCT in the Monte Carlo method may further improve the fidelity of the estimated OCT signal [36]. Efforts are currently underway to expand the implementation of the Monte Carlo to situations where significant spatial variation in the scattering phase function *p*(*θ, φ*) is expected [37], as is the case in glandular mucosa. Finally, material inhomogeneities inside each tissue layer can be accounted for using models of epithelial morphogenesis [38], although simultaneous resolution of deformations due to small-scale and tissue-scale stresses requires a multiscale treatment that is yet to be developed. The mesh-based Monte Carlo method facilitates the integration of biomechanical modeling in the existing data generation pipeline.

Among the priorities for future work is the resolution of inter-patient variability focusing on known biases in the target patient population. Candidates for BE surveillance present with tissue damage from long-term chronic reflux disease and epithelial alterations due to persistent esophagitis. Estimates of the optical properties in the control population may therefore require further adjustments for deviations from the healthy stratified squamous epithelium. Hiatial hernias were present in the majority of the patients in our cohort, requiring further examination of the differences between the esophageal and gastric mucosa, and those between the gastric and Barrett’s mucosa. To a first approximation, we aggregated all tissue states prior to the onset of glandular morphogenesis into one label in the present work. Similarly, we aggregated the states corresponding to glandular mucosae of the gastric and esophageal phenotypes. We expect the expansion of the set of labels considered in the classification problem and inverse modeling of optical properties to inform data generation to improve the performance of our pipeline.

## V. Conclusions

Computational approaches to data augmentation represent a promising approach to overcoming data scarcity in the application of deep learning to diagnostic surveillance of rare conditions via tissue imaging. Physically-motivated computations that rely on clinical knowledge and mechanistic understanding of disease may provide an advantage over limited clinical datasets in data-poor settings. We demonstrated the utility of this approach for a proof-of-concept application to the classification of esophageal OCT.

## VI. Acknowledgements

We would like to thank Jianhua Lee, Brianna Lauren, Aaron Oh, and Lindsay Kumble for their help with the curation and review of EHR data, Zhong Wang of the Digital and Computational Pathology Laboratory at CUIMC for help with digitization of pathology slides, and Nicholas Giangreco for help with handling of OCT data. Fruitful conversations with NinePoint Medical (Bedford, MA) regarding the OCT data are appreciated. This work was supported by the National Institutes of Health via the grants U01 CA 199336 and R01 CA 247790. The authors declare no conflicts of interest.

## VII. Supplementary Information

**TABLE II.** (SUPPLEMENT) Glossary of terms extracted from pathology reports and their correspondence to reduced labels indicating the underlying microanatomy and disease stage. The final label describes the annotation in the absence of any other indication.

| Keyword | Abbrev. | Label |
| --- | --- | --- |
| esophagitis | ESGTS | STRATIFIED EPITHELIUM |
| inflammation | INF | STRATIFIED EPITHELIUM |
| reflux esophagitis | RFLX_ESGTS | STRATIFIED EPITHELIUM |
| reactive features | R | STRATIFIED EPITHELIUM |
| intraepithelial eosinophil | INTR_EOSIN | STRATIFIED EPITHELIUM |
| increased eosinophils | INCR_EOSIN | STRATIFIED EPITHELIUM |
| squamous mucosa | SQ_MUCOSA | STRATIFIED EPITHELIUM |
| mild reactive features | M_R | STRATIFIED EPITHELIUM |
| reflux | RFLX_INDICATED | STRATIFIED EPITHELIUM |
| columnar epithelium | COLE | METAPLASIA |
| consistent with Barrett's esophagus | BE | METAPLASIA |
| rare goblet cells | RGC | METAPLASIA |
| goblet cells | GC | METAPLASIA |
| positive for intestinal metaplasia | IM | METAPLASIA |
| squamocolumnar mucosa | SCOLE | METAPLASIA |
| reflux carditis | RFLX_CDTS | STOMACH |
| cardia-type mucosa | CM | STOMACH |
| gastric cardia-type mucosa | CM | STOMACH |
| cardiac-type mucosa | CM | STOMACH |
| carido-oxtyntic-type mucosa | COM | STOMACH |
| gastric cardio-oxtyntic mucosa | COM | STOMACH |
| gastric oxyntic-type mucosa | COM | STOMACH |
| gastric oxyntic mucosa | COM | STOMACH |
| cardio-fundic type mucosa | CFM | STOMACH |
| fundic-type mucosa | FM | STOMACH |
| low-grade dysplasia | LGD | DYSPLASIA |
| high-grade dysplasia | HGD | DYSPLASIA |
| intramucosal adenocarcinoma | IMC | CANCER |
| adenocarcinoma | AC | CANCER |
| high-grade squamous dysplasia | SHGD | OTHER |
| low-grade squamous dysplasia | SLGD | OTHER |

**Fig. 3.**
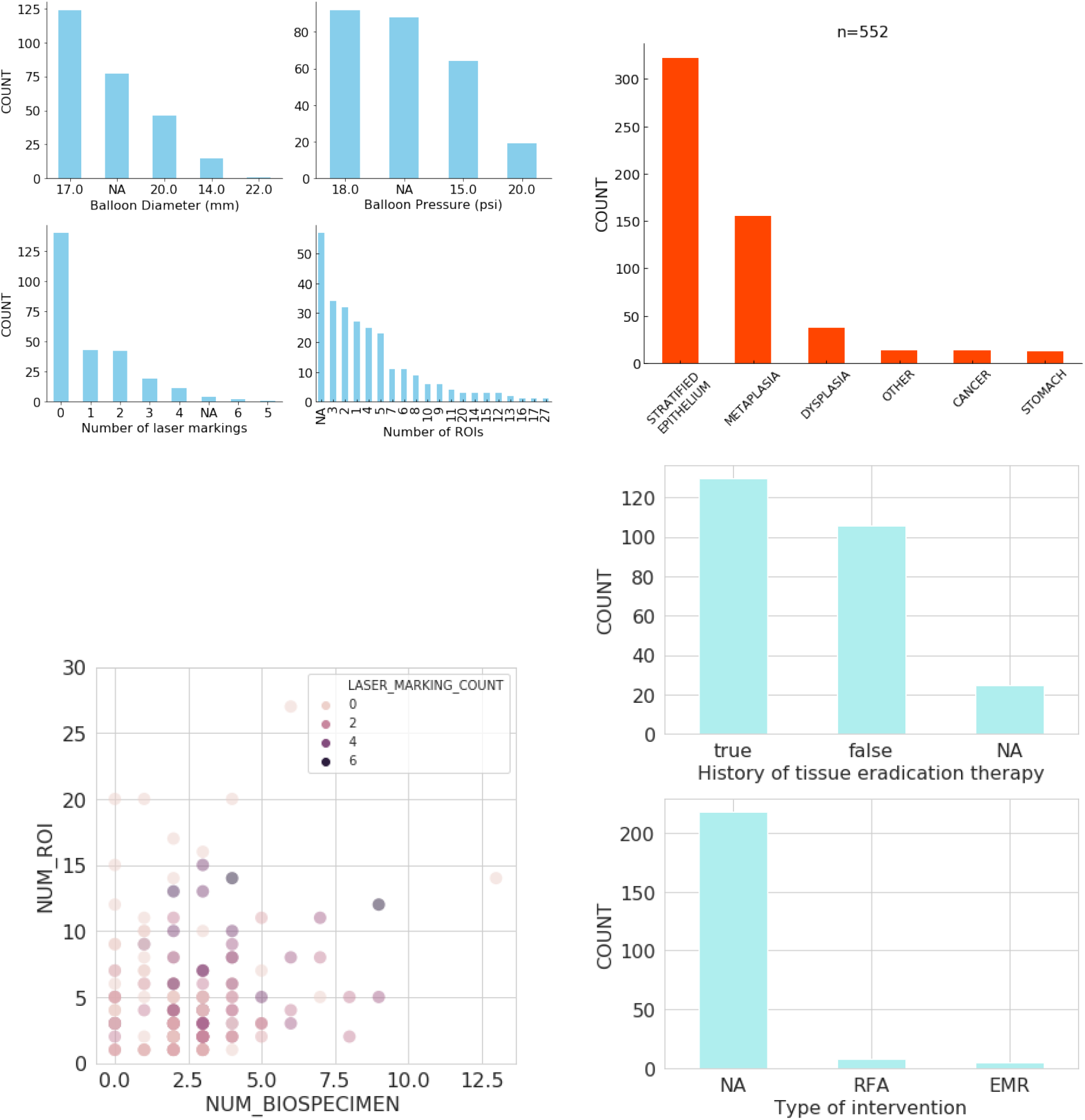
(SUPPLEMENT) Examples of missingness and heterogeneity in the imaging and pathology datasets: (top left: A) The number of ROIs and laser-marked regions vary among scans. Different balloon catheter diameters and pressures had been used during the period of data collection. (top right: B) Annotated biospecimens were skewed toward controls (stratified epithelium = `NORMAL`). (bottom left: C) Tissue sampling is guided by the endoscopist’s on-the-fly assessment of the risk of progression, and the extent of *ex vivo* confirmation needed on a patient-by-patient basis. (bottom right: D) Patients present at different stages of diagnosis and treatment, some for follow-up after interventions like tissue eradication therapy, and different treatments vary in terms of their impact on existing tissue; e.g. radiofrequency ablation (RFA) may remove large segments of the epithelium while endoscopic mucosal resection (EMR) is a localized intervention.

